# A longitudinal evaluation of localised chronic *Pseudomonas aeruginosa* infection in cystic fibrosis rat models

**DOI:** 10.1101/2025.04.11.647899

**Authors:** Nicole Reyne, Bernadette Boog, Patricia Cmielewski, Alexandra McCarron, Ronan Smith, Nathan Rout-Pitt, Nina Eikelis, Kris Nilsen, John Finnie, Jennie Louise, David Parsons, Martin Donnelley

**Affiliations:** Robinson Research Institute, University of Adelaide, Adelaide, South Australia, Australia; Adelaide Medical School, University of Adelaide, Adelaide, South Australia, Australia; Respiratory and Sleep Medicine, Women’s and Children’s Hospital, Adelaide, South Australia, Australia; 4DMedical, Melbourne, Victoria, Australia; Biostatistics Unit, South Australian Health and Medical Research Institute, Adelaide, South Australia, Australia

**Keywords:** *Pseudomonas aeruginosa*, lung infection, cystic fibrosis, animal model

## Abstract

Recurrent bacterial infections with *Pseudomonas aeruginosa* result in chronic airway inflammation, lung damage and eventual respiratory failure, and are the major cause of morbidity and mortality in people with cystic fibrosis (CF). Animal models are essential for understanding disease progression and assessing potential treatments in the presence of infection. Previously reported *P. aeruginosa* lung infection rodent models for CF research have some weakness, including acute infection rather than chronic, associated mortality, use of laboratory strains of *P. aeruginosa* and the use of non-CF rodents. The aim of this study was to create a localised *P. aeruginosa* infection in wildtype and two CF rat models, by delivering bacteria embedded agar beads using a miniature bronchoscope. The resulting infection was well tolerated by all animals of all genotypes with no mortality associated with the procedure or infection. Histologically the affected regions were localised to the right lung, with bronchopneumonia present. Bacteria persisted for 9 weeks (63 days) in all genotypes, with lung function changes observed by day 63 of the infection.

## Introduction

*Pseudomonas aeruginosa* (*P. aeruginosa*) is a major pulmonary bacterial pathogen in people with cystic fibrosis (CF), causing respiratory infections in approximately 80% of individuals [1]. *P. aeruginosa* is an aerobic, gram-negative, rod-shaped bacterium found in various environments. While it can cause opportunistic infections such as pneumonia, wound infections, and urinary tract infections in humans, it rarely affects healthy lungs [2]. Individuals with CF who have compromised lung defence mechanisms are particularly vulnerable to *P. aeruginosa* infections. This microorganism demonstrates remarkable adaptability, undergoing genetic, physiological, and morphological alterations within the CF lung environment. One adaptation is the over-production of alginate, forming thick biofilms that protects the bacteria from phagocytosis and antibiotic therapy [3]. Initial infections may respond to aggressive antibiotic therapy, but once *P. aeruginosa* becomes established in the airways of people with CF, it is nearly impossible to eradicate, leading to chronic infections [4].

To develop effective lung-directed treatments for CF, it is crucial to test them under conditions that closely recapitulate the CF lung environment. CF animal models vary in degrees of CF pathophysiology, often exhibiting gut obstructions, reproductive defects and nasal bioelectrical defects [5]. However, few models develop the chronic lung disease characterised by pathogen colonisation. To better mimic CF lung disease, methods of administering P. *aeruginosa* to the lungs in various forms have been developed. There are various factors to consider when creating an *P. aeruginosa* infection, including *P. aeruginosa* strain, delivery technique, length of infection and the choice of rodent strain [6].

Animal models have been developed to represent either the acute or chronic stage of the *P. aeruginosa* infection. Acute models generally involve the delivery of planktonic *P. aeruginosa*, which can lead to either rapid clearance or severe septic infection [7-9]. It is important to develop animal models that replicate the chronic, persistent infections observed in people with CF. One common approach involves delivering agar beads containing or mixed with *P. aeruginosa* to the airways [10]. This technique retains the bacteria in the lungs, creating a persistent infection more like those observed in people with CF.

Traditionally, the delivery of bacteria into the lungs of rodents has been performed using an endotracheal tube placed via a tracheostomy, carrying a risk of mortality [11]. Our group has developed and validated bronchoscopic procedures in rat lungs by adapting a small sialendoscope normally used in humans [12, 13]. This technique allows precise and accurate delivery to rat lungs without the need for surgical incision, significantly reducing mortality. The procedure is quick (less than 1 minute), can be performed using inhaled anaesthesia, and enables targeted delivery of agar beads to a specific region of the lung [14]. This approach allows rats to maintain adequate health and facilitate the assessment of treatments or interventions. It may also better reflect the patchy lung disease seen in people with CF [15].

In this study, we employed a miniature bronchoscope to create a localised chronic *P. aeruginosa* infection in the right lung of wildtype and two CF rat strains (knockout and *Phe508del*) using bacteria-embedded agar beads. We analysed the inflammatory response in the bronchoalveolar lavage (BAL), confirmed the stability of bacteria load at three weeks and out to nine weeks, explored the histological changes, and assessed the effect on lung function using flexiVent lung mechanics and X-ray Velocimetry ventilation measurements.

## Methods

### Animals

This project was conducted under the approval of the University of Adelaide (M-2024-078) and South Australian Health and Medical Research Institute (SAM424.19) Animal Ethics Committees, and performed in accordance with ARRIVE guidelines [16]. Male and female (average 16 weeks) CFTR knockout (n = 33), *Phe508del* (n = 34) and wildtype (n = 38) littermate control rats were used [17]. Rats were maintained in individually ventilated cages (IVC) with a 12-h light/dark cycle.

### Bacterial strain

All studies used *P. aeruginosa* 20844 muc, a polymyxin susceptible strain that was collected from a CF patient and supplied by Monash University. Bacteria were maintained in glycerol stocks and stored at -80°C.

### *P. aeruginosa* embedded agar beads

Agar beads were prepared by a modification of existing methods [18]. The *P. aeruginosa* 20844 muc was cultured overnight in 30 mL of Luria Bertani (LB) broth in a shaking incubator at 280 rpm at 37°C. Bacteria (10^9^ CFU) were centrifuged at 5,300 x g (rcf) for 10 minutes and resuspended in 6.25 mL of LB broth. 25 ml of 2% agarose (A4679, Sigma) and 80 mL of mineral oil were heated separately to 53°C while stirring at a medium speed. The bacteria were added to the agarose and briefly mixed. Then, 2.5 mL of the bacteria/agarose mixture was syringed into the mineral oil while stirred rapidly and cooled for 10 minutes. Deoxycholic acid (DCA) was used to wash the agar beads twice by centrifuging at 690 x g (rcf) for 10 minutes and removing the top layer of oil each time. This was followed by three washes in phosphate buffered saline (PBS). The bacteria-laden agar beads were resuspended in an equal volume of PBS to form a *P. aeruginosa* agar bead slurry, and the number of bacteria present was determined by plating 10-fold serial dilutions on LB agar plates. A total of nine agar bead preparations were used for this study, with an average 2.4 × 10^6^ CFU/ml and bead size of 50 - 550 μm.

### Rat model of localised *P. aeruginosa* infection

To generate a localised lung infection model, rats were deeply anesthetised using 3% isoflurane and suspended on an intubation stand by their incisors. A Hamilton syringe (250 μl) was attached to the miniature bronchoscope (Storz, rigid endoscope, Model 11582 A, working channel size 350 μm) via an ∼30 cm tube, to draw up 50 μl (1.2 × 10^5^ CFU) of the *P. aeruginosa* bead slurry and a 50 μl of air chaser into the working channel. The bacteria laden beads were delivered to the top of the right main bronchus [12]. The rats were then allowed to recover and were monitored and weighed daily.

Assessments were conducted in separate groups of rats on days 7-, 14-, 21- and 63-days following infection, after which the rats were humanely killed (no repeated measures were performed). The rats were divided into two groups: **group 1** was used for lung function assessment (see XV imaging and flexiVent mechanics, below) followed by assessment of bacteria colony forming units (CFU) to determine the bacterial load, and **group 2** was used for lung function assessment followed by bronchoalveolar lavage (BAL) fluid analysis (day 63 not assessed) and lung histology.

### X-ray Velocimetry (XV) imaging

Rats were XV imaged as previously described [14, 19]. Briefly, rats were anaesthetised with medetomidine (0.4 mg/kg) and ketamine (60 mg/kg) by intraperitoneal injection. Once anaesthetised, the rats were prepared by performing a tracheostomy and cannulation with a cut-down endotracheal tube (ET; 14 Ga BD Insyte plastic cannula bevel cut to 15 mm length), and then positioned upright (head-high) into a animal holder and placed onto the rotation stage of the Permetium preclinical scanner (4DMedical, Australia). The rotation stage was 1700 mm to detector distance. Rats were ventilated using a pressure controlled small animal ventilator (4DMedical Accuvent 200, Australia) set to a peak inspiratory capacity if 14 cmH_2_O, positive end-expiratory pressure of 2 cmH_2_O, and ventilated at 93.75 breaths/min (192 ms inspiration and 448 ms expiration, I:E ratio 3:7). A 4D XV scan was acquired at a frame rate of 15.625 Hz, with 10 images per breath and 600 projections per phase point. The scan time was ∼ 6 min.

Specific ventilation was only assessed in the inspiratory phase of the breath. All XV data was analysed by 4D Medical using a proprietary algorithm that calculates mean specific ventilation (MSV), ventilation heterogeneity (VH), ventilation defect percentage (VDP) and tidal volume. Further analysis of the lung was done using a custom Python script that bisected the lungs into right and left halves [20].

### Assessment of respiratory mechanics by flexiVent

Following XV imaging, lung function assessments were performed using a flexiVent FX small animal ventilator fitted with a FX4 rat module (SCIREQ, Montreal, Canada). The rats were placed into a supine position and the ET was connected to the flexiVent and measurements (using the Deep Inflation, SnapShot-90, Quick Prime-3, and PVs-P perturbations) were performed as previously described [19]. Three measurements for each parameter were made for each rat and these were averaged. Rats were then humanely killed by i.p overdose of lethabarb (150 mg/kg, Virbac, Australia).

### Bacteriology

The right lung of group 1 was excised and mechanically homogenised (Omni, TH220) in 3 ml PBS and serially diluted (1:10 - 1:10,00). Diluted lung homogenate (100 μl) was plated on LB agar for incubation at 37°C overnight and colonies were manually counted.

### Bronchoalveolar lavage

Bronchoalveolar lavage (BAL) was performed on the right lung of group 2 (excluding day 63), by exposing the lungs and tying off the left lobe. The right lobe was lavaged with 1.5 mL of warmed phosphate-buffered saline (PBS), then 500 mM ethylenediaminetetraacetic acid (EDTA) and 100 mM phenylmethylsulfonyl fluoride (PMSF) were added to the BAL fluid. Lavage was cytospun onto slides and stained using Giemsa. The number of macrophages, lymphocytes and neutrophils were quantified by randomly selecting three fields of view and a counting a minimum total of 300 cells per animal using a Nikon Eclipse E400 microscope (Tokyo, Japan).

### Histology

Lungs from group 2 were fixed in 10% neutral phosphate-buffered formalin, embedded in paraffin and sectioned at 5 μm. Sections were stained with Hematoxylin and Eosin and duplicate sections were stained with Alcian blue/Periodic acid Schiff to detect the presence of mucus. Images were examined by a pathologist and captured on a Nikon Eclipse E400 microscope with DS-Fi2-U3 camera and Nikon NIS-elements D software version 5.42.08 (Tokyo, Japan).

Immunohistochemistry was used to locate *P. aeruginosa* in the lungs of infected rats. Slides were deparaffinised and rehydrated using standard histological procedures. Antigen retrieval was performed using a 10 mM sodium citrate buffer (pH 6) for 20 minutes. A protein block was applied for 30 minutes using 2% bovine serum albumin (BSA) and 0.1% Triton x-100. Sections were incubated overnight at 4°C with rabbit polyclonal to *pseudomonas* (1:600) (ab68538, Abcam) in 2% BSA and 0.1% Triton. Sections were rinsed and incubated in donkey anti-rabbit (Alexa Fluor 558) (IgG) (1:500) (ab175470, Abcam) made in 2% BSA and 0.1% Triton for one hour at room temperature. The slides were washed, counterstained with DAPI for approximately 10 minutes, washed and mounted with Prolong Diamond antifade mounting media (#P36961, Life Technologies, USA). Slides were visualised for the presence of *P. aeruginosa* using a Zeiss LSM980 super-resolution confocal microscope with airyscan 2 (Germany).

### Statistics

Statistical analyses were performed in R version 4.4.1 [21]. To assess differences in the flexiVent and XV parameters from baseline to each infection assessment point, a linear regression model was fitted to each parameter using the *lm* function. This model contained fixed effects of genotype, weight and infection timepoint. Pairwise comparisons for each fitted model were performed using the *emmeans* package [22]. Results are presented graphically as estimated marginal means and 95% confidence intervals.

## Results

### Animal health

The delivery of bacteria embedded agar beads was well tolerated, with no mortality from the delivery using the miniature bronchoscope. The rats were assessed daily until 21 days, then weekly, throughout the study for weight, health and well-being using a clinical record sheet, with no significant adverse effects on the animals’ health observed. Further, there were no deaths associated with the infection for the duration of study. A small transient weight loss was observed in the first 3 days following infection, irrespective of the animal genotype. All rats returned to pre-procedure weight and then continued to gain over the remainder of the study period.

### Bacterial load

The average bacterial count recovered from the right lung homogenate of each genotype (wildtype: 1.9 × 10^6^, *Phe508del*: 8.6 × 10^5^, knockout: 4.6 × 10^6^ CFU/lung) was higher than the initial inoculum (1.2 × 10^5^ CFU on average) at day 7 (Figure 1A). There were no significant differences in the CFU counts between genotypes at any time point. By day 63, wildtype and *Phe508del* rats had a lower mean CFU count lower than the initial inoculum (wildtype: 5.9 × 10^4^ CFU, *Phe508del*: 3.7 × 10^4^, knockout 6.5 × 10^5^ CFU/lung).

**Figure 1:**
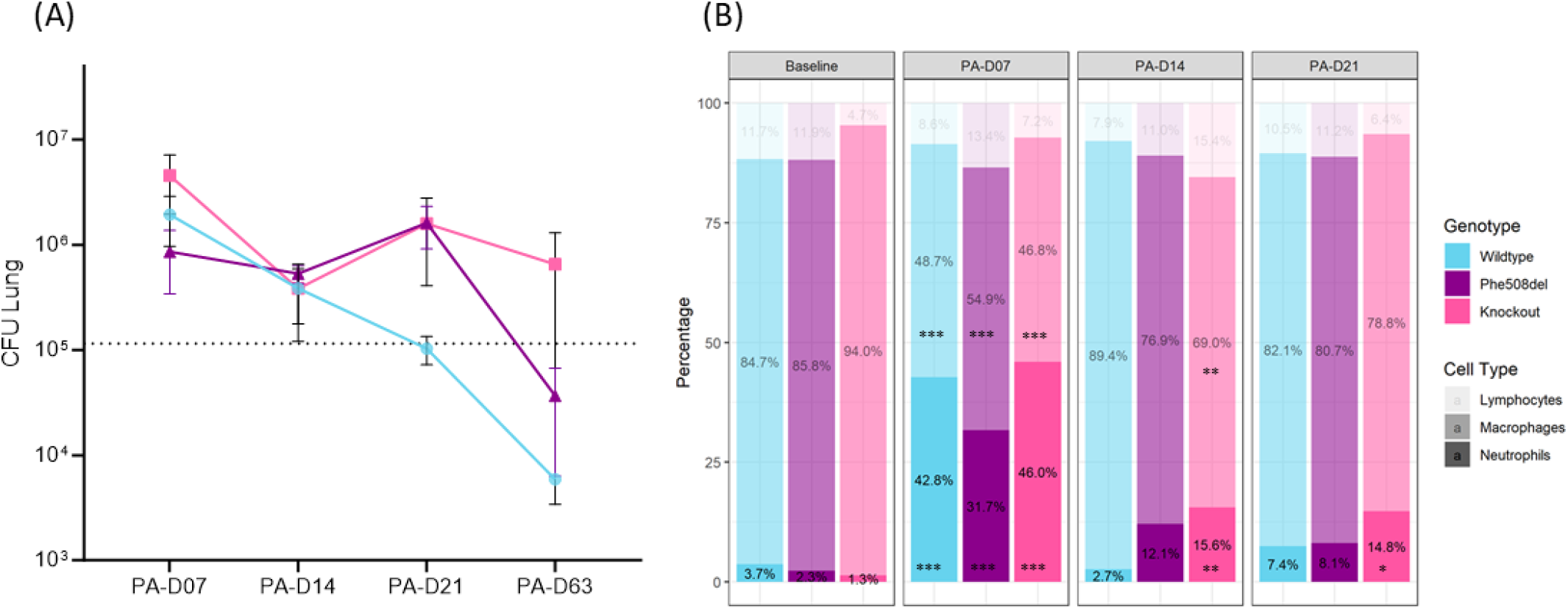
(A) Bacterial load in lung homogenates and (B) bronchoalveolar lavage (BAL) cell proportions (%) following *P. aeruginosa* agar bead inoculation in wildtype, *Phe508del* and knockout rats. (A) Log CFU/gram of lung at days 7, 14, 21 and 63 post infection (n = 2 - 6, dash line = average CFU in 50 μl delivered). (B) Lymphocyte, macrophage and neutrophil proportion in BAL at days 7, 14, and 21 post infection. (n = 3 animals/genotype, linear regression model, estimated marginal means and 95% CIs, * *p*<0.05, ** *p*<0.01, ** *p*<0.001 within genotypes compared to baseline). Key on graph: PA-D07 = 7 days post *P. aeruginosa* bead delivery; PA-D14 = 14 days post *P. aeruginosa* bead delivery; PA-D21 = 21 days post *P. aeruginosa* bead delivery, and PA-D63 = 63 days post *P. aeruginosa* bead delivery.

### Bronchoalveolar lavage analysis

At day 7 all genotypes exhibited an increase in the percentage of neutrophils and a corresponding decrease in the percentage of macrophages post *P. aeruginosa* inoculation (Figure 1B). Neutrophils remained raised in the knockout rats and were significantly different from baseline at day 21, while in wildtype and *Phe508del* rats there was no difference at day 21 compared to baseline, with neutrophils returning to baseline levels. BAL was not assessed on day 63.

### Histopathology

Histopathological changes, of varying severity, were localised to the bottom right lung in all rats and at all timepoints (Figure 3). The two principal morphological changes were an acute suppurative bronchopneumonia associated with the bacteria-laden agar beads and a lymphocytic vasculitis. In all rats these pathological features in affected lung lobes were notably multifocal in distribution, with large intervening areas of unaffected parenchyma.

**Figure 3:**
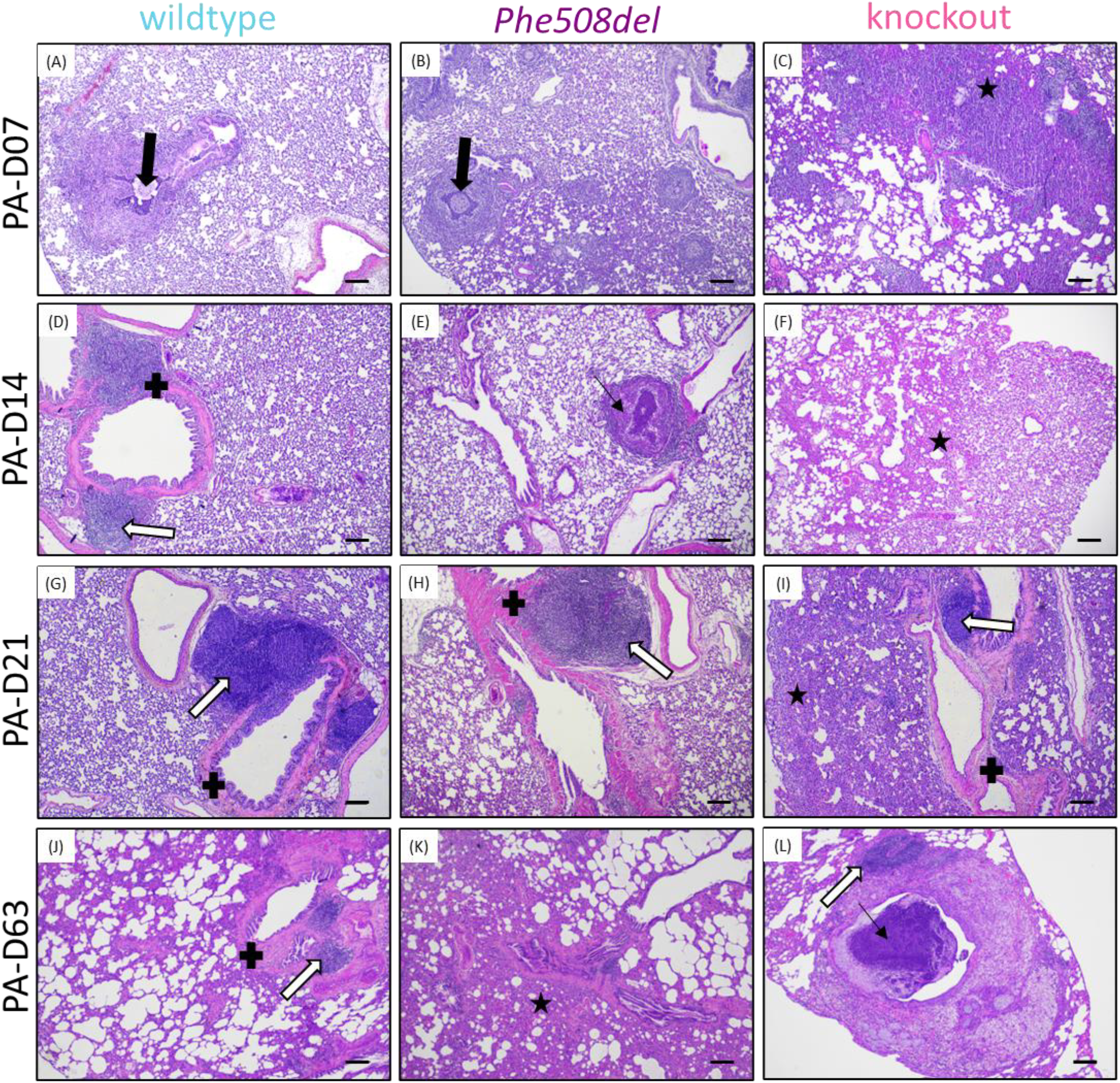
Representative haematoxylin and eosin images from 7-, 14-, 21-, and 63-days post-infection in (A, D, G, J) wildtype, (B, E, H, K) *Phe508del* and (C, F, I, L) knockout rats. Thick black arrow indicates agar bead, thin black arrow neutrophils in airway, white arrow shows areas of BALT, black star alveolar thickening, and black cross peribronchial thickening. (Scale bar 200 μm).

The agar beads containing bacteria in bronchial and bronchiolar lumina were surrounded by abundant neutrophils (Figure 3), with acute inflammatory cell infiltrates in the mucosa. The bronchopneumonia was often necrotizing, with the lumina being filled with obstructing, desquamated, necrotic cellular debris, which was admixed with neutrophils and bacteria. The acute bronchopneumonia sometimes extended into contiguous alveolar spaces, with aggregate neutrophils and macrophages. Agar beads disintegrated over time, but the acute bronchitis/bronchiolitis persisted and the number of macrophages in, and around, affected airways progressively increased with time. There was frequently a severe, often necrotizing, lymphocytic vasculitis, with mural lymphocytic infiltration and often attenuation, or even segmental absence, of intact vessel wall; in a few vessels, there was fibrinoid necrosis. These blood vessels were surrounded by large cuffs of lymphocytes, and fewer plasma cells (lymphoblastic cuffing). The alveolar interstitium adjacent to the inflamed bronchi and bronchioles was often thickened by varying numbers of lymphocytes, macrophages and neutrophils. Peribronchial thickening was observed across all genotypes, with the degree of thickening varying throughout the course of infection. There was commonly bronchial associated lymphoid tissue (BALT) hyperplasia (Figure 3D, G, H, I, J, L), with lymphoid follicles showing active germinal centres. Some bronchi and large bronchioles showed marked goblet cell hyperplasia (Figure 4), with extruded mucus often overlying the epithelial surface or plugging the airway lumina (Figure 4). Knockout rats exhibited widespread alveolar thickening at all stages of the infection, while wildtype and *Phe508del* rats demonstrated focal thickening with surrounding alveolar spaces remaining clear.

**Figure 4:**
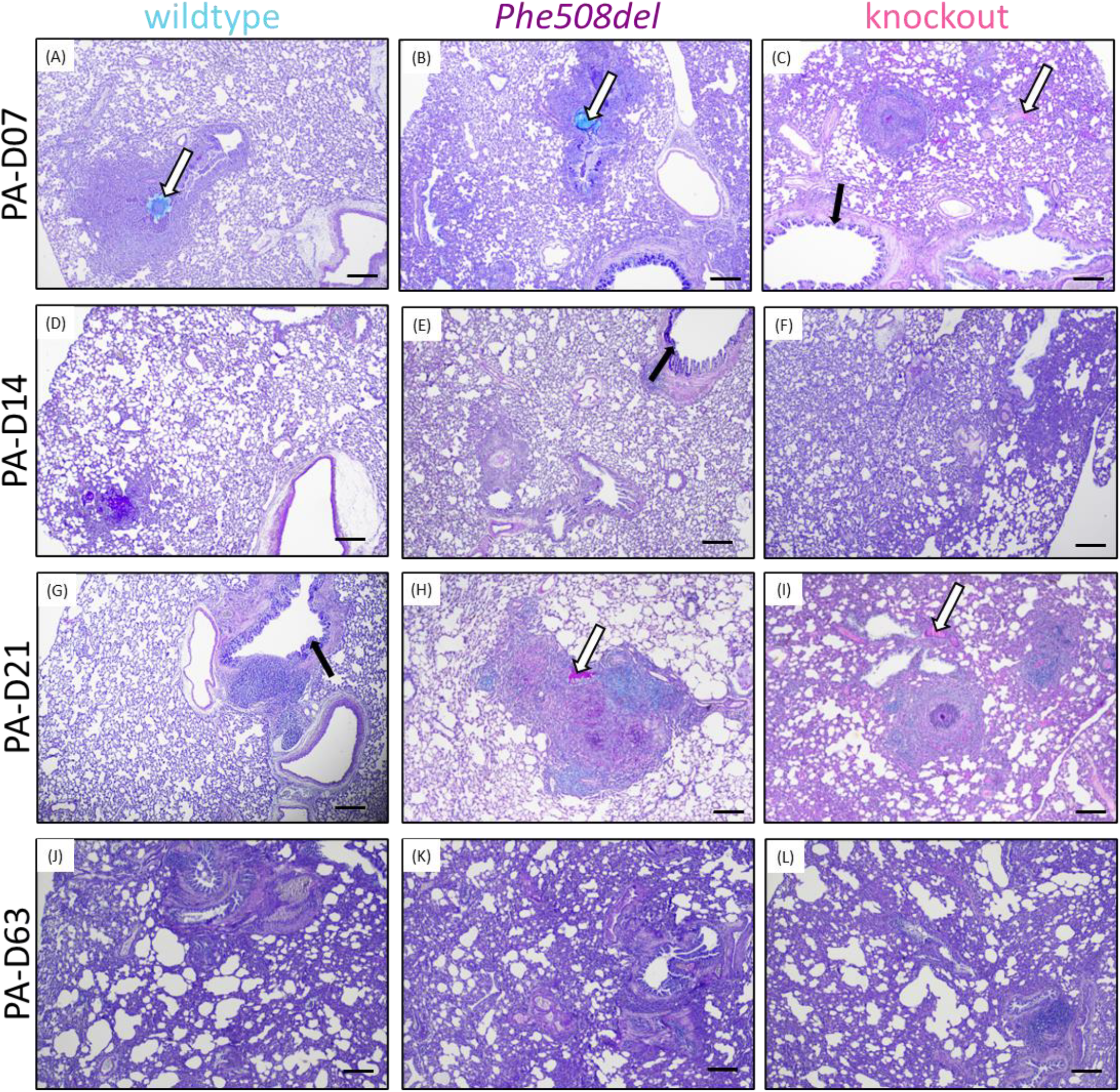
Representational Alcian Blue-Periodic Acid-Schiff images from 7-, 14-, 21- and 63-days post-infection in (A, D, G, J) wildtype, (B, E, H, K) *Phe508del* and (C, F, I, L) knockout rats. Black arrows indicate goblet cell hyperplasia and white arrows mucus plugging. (Scale 200 μm).

At day 63 of infection, histological analysis revealed consistent and severe features, with the right lung exhibiting more pronounced changes than the left lung. The primary finding was chronic, multifocal interstitial pneumonia, characterised by infiltration of the alveolar interstitium by macrophages and lymphocytes, alongside varying degrees of fibrosis and emphysema (Figure 3J-L). Alveolar emphysema was evident as enlargement and coalescence of airspaces, caused by destruction and loss of alveolar septa. This change was more pronounced in the subpleural regions of the right lung. Additionally, alveolar macrophages, many of which were foamy, were present within the alveolar spaces and occasionally aggregated into epithelioid granulomas. In the right lungs, luminal accumulations of extruded mucus were observed in the bronchi and bronchioles, with some extending into alveoli when the mucus load was abundant. When mucus accumulation was particularly copious, there was goblet cell hyperplasia within the airways.

Immunohistochemistry was used to verify the presence of *P. aeruginosa* in the lungs (Figure 5). At early stages of the infection (day 7), *P. aeruginosa* was detected within the agar beads, and at later stages (day 21) the *P. aeruginosa* was more diffuse across the lung tissue.

**Figure 5:**
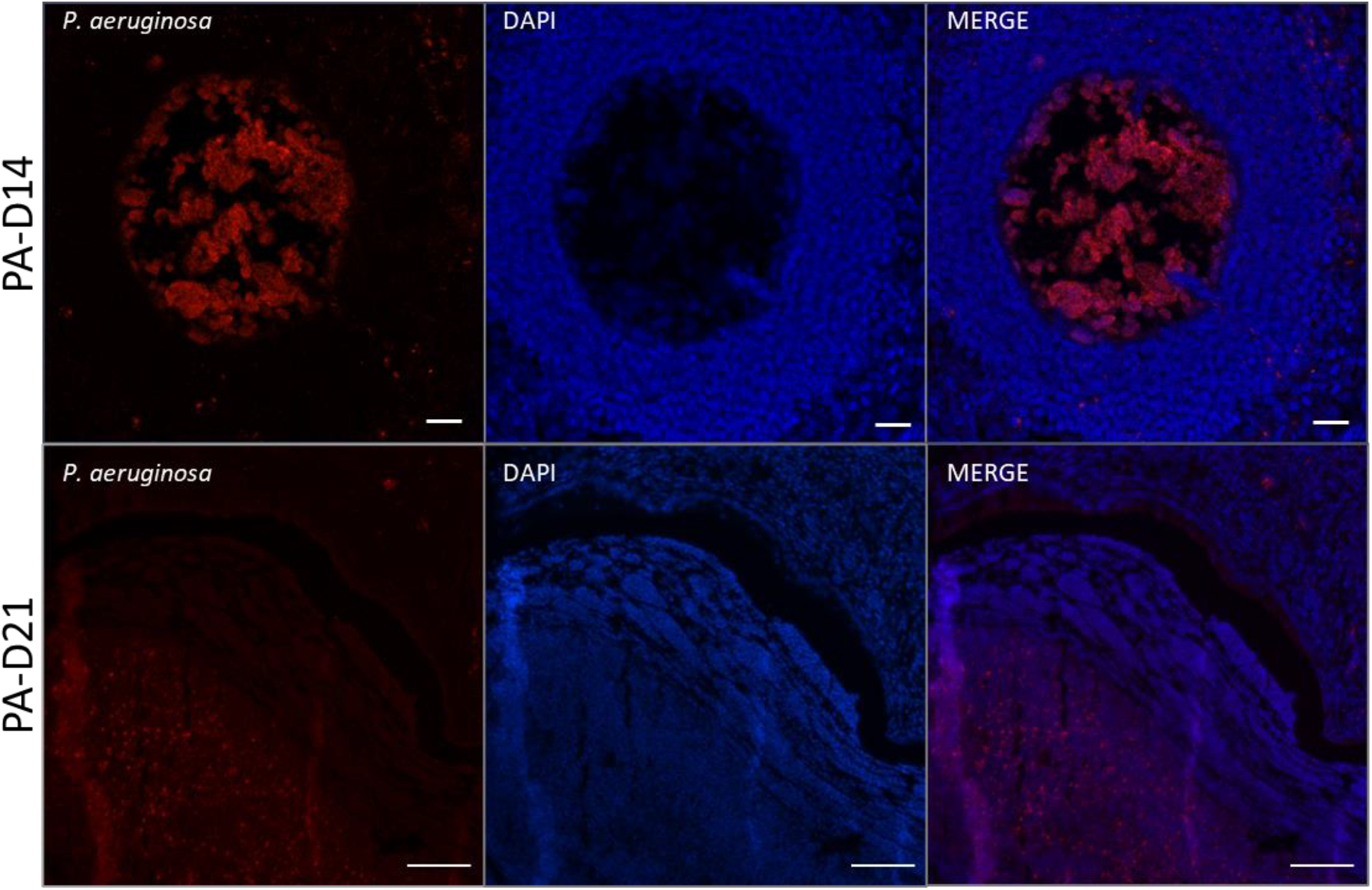
Immunohistochemistry detection of *P. aeruginosa* in the lung at 7- and 21-days post infection. Bacteria were localised using an anti-*P. aeruginosa* (red) antibody with DAPI counterstain (blue). (7-day image from *Phe508del* and 21-day image from knockout rat)

### FlexiVent respiratory mechanics

In the wildtype rats there was an increase in IC_norm_ with day 63 of the infection being significantly increased from all other timepoints. While *Phe508del* rats exhibited an increase from baseline to days 7, 14 and 63. At day 63 the knockout rats had a significant increase in IC_norm_ compared to baseline (Figure 6A).

**Figure 6:**
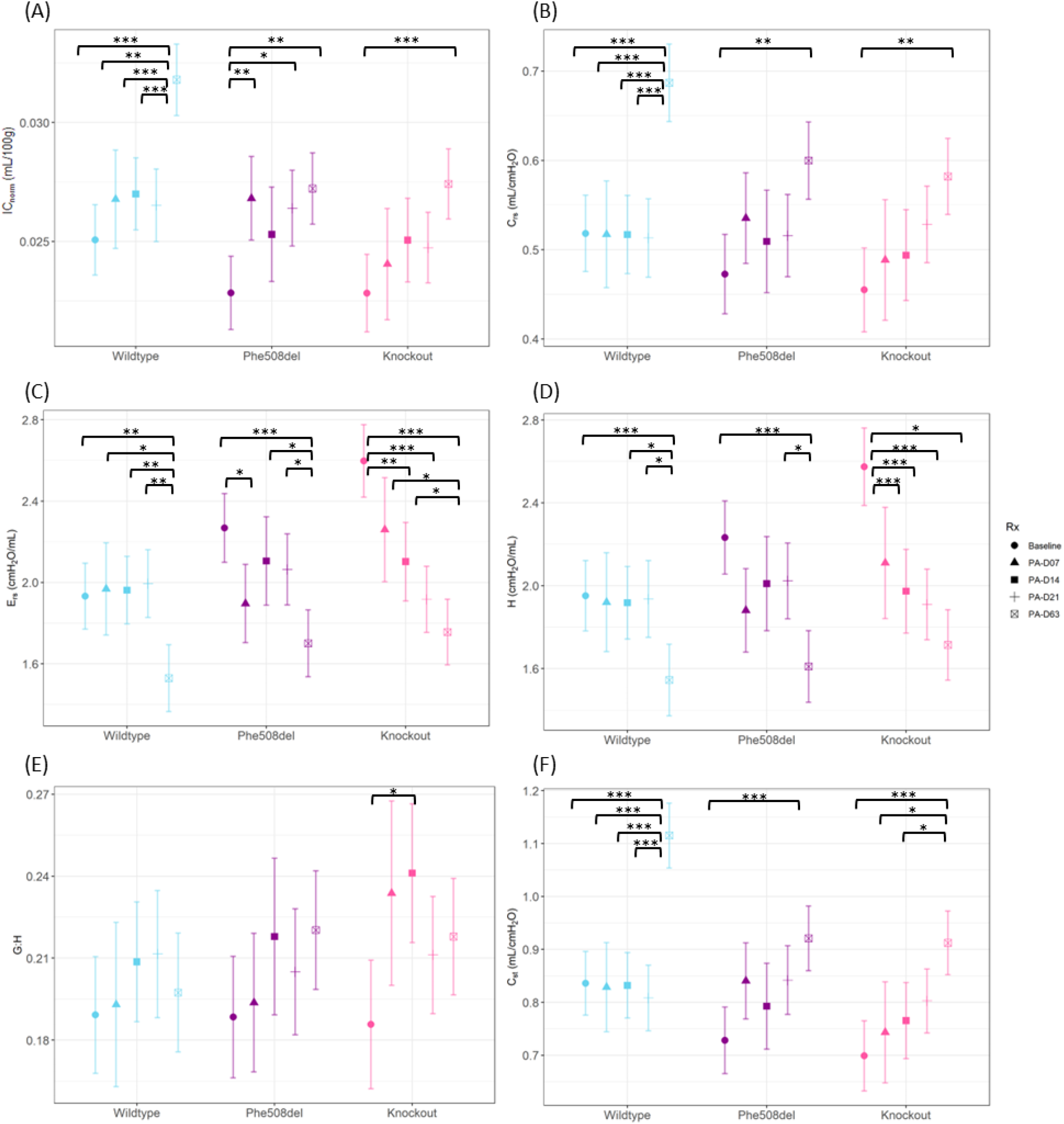
FlexiVent assessments of the respiratory system in wildtype, Phe508del and knockout rats at baseline and days 7, 14, 21 and 63 post local *P. aeruginosa* infection. Deep inflation was used to produce the (A) inspiratory capacity (IC_norm_). The single frequency forced oscillation measured the (B) total respiratory system compliance (C_rs._) and (C) total respiratory system elastance (E_rs_). The broadband forced oscillation measured the (D) tissue elastance (H) and (E) tissue hysterisivity (G:H) of the peripheral lung compartment. The pressure volume loop (G) was used to determine the static compliance (F). (n = 3-7 animals/genotype, linear regression model, estimated marginal means and 95% CIs, * *p*<0.05,** *p*<0.01 \*\*\**p*<0.001). Key on graph: Baseline = no PA infection, PA-D07 = 7 days post *P. aeruginosa* bead delivery; PA-D14 = 14 days post *P. aeruginosa* bead delivery; PA-D21 = 21 days post *P. aeruginosa* bead delivery; and PA-D63 = 63 days post *P. aeruginosa* bead delivery.

The single frequency forced oscillation (single compartment model) analysis revealed decreased total respiratory system elastance (E_rs_) in all genotypes from baseline to day 63 post-infection (Figure 6C). Total respiratory system compliance (C_rs_) (Figure 6B) was significantly increased in all genotypes from baseline to day 63.

The broadband forced oscillation (constant phase model) showed no difference in Newtonian (central airway) resistance (R_n_) at any time point or genotype. Tissue elastance (H) was significantly different between the genotypes at baseline, and all genotypes also displayed a decrease in elastance from baseline to day 63 post infection. Tissue damping (G) was significantly lower in wildtype rats compared to *Phe508del* and knockout rats at baseline. Knockout rats showed a decrease in tissue damping over the infection period (Figure 6D-E).

Average pressure-volume loops constructed from mean data were used to determine the static compliance (C_st_). Static compliance was higher in wildtype rats compared to *Phe508del* and knockout rats at baseline, and there was a significant increase in static compliance from baseline to day 63 in all genotypes (Figure 6F).

### X-ray Velocimetry (XV) imaging

There were no changes in the MSV and VDP in any genotype over infection period (Figure 7A-B). The knockout rats demonstrated a significant difference from day 21 to day 63 in VH (Figure 7C)

**Figure 7:**
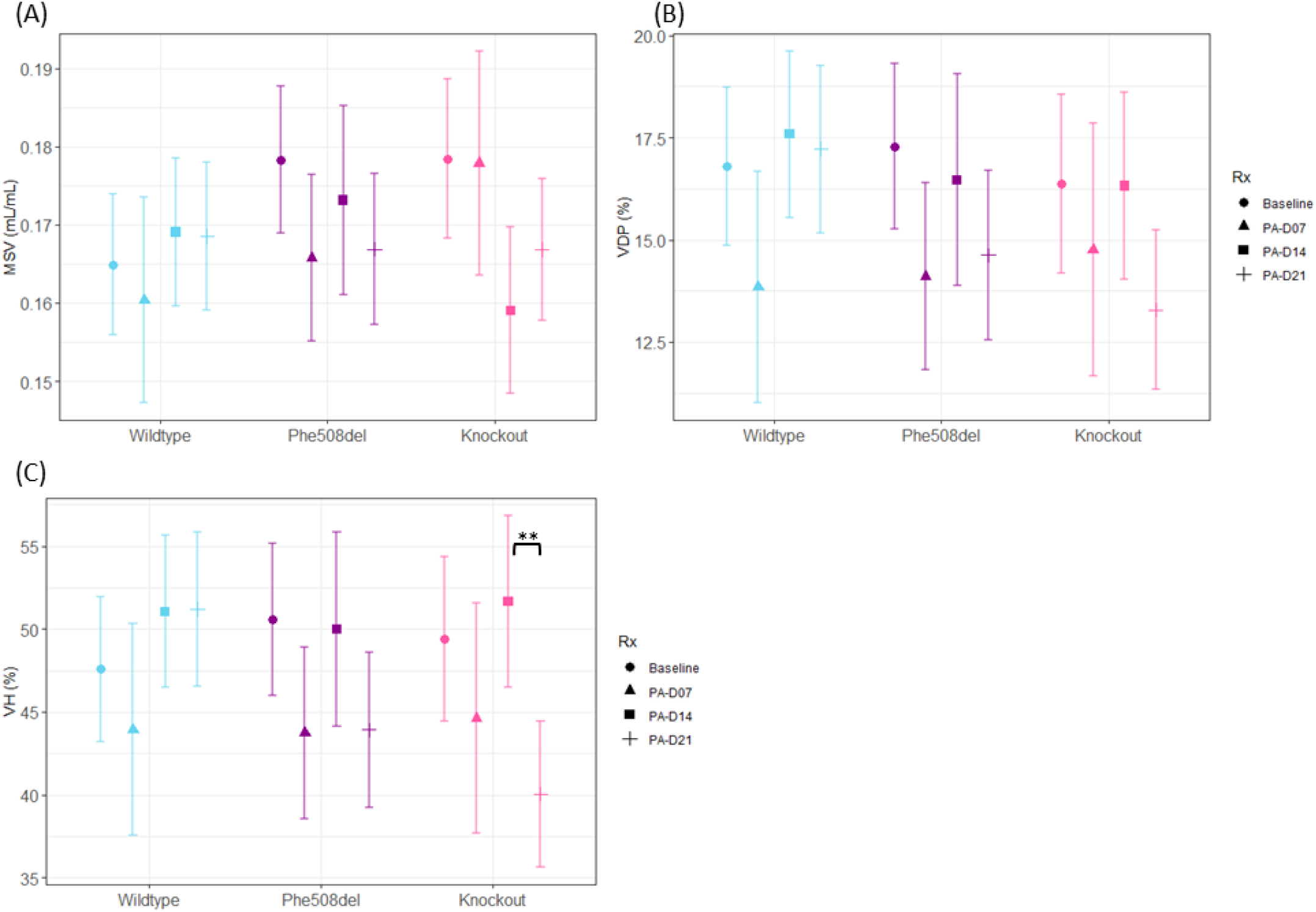
X-ray velocimetry parameters in wildtype, Phe508del and knockout rats at baseline and days 7, 14, 21 and 63 post local *P. aeruginosa* infection. (A) Mean specific ventilation, (B) ventilation defect percentage and (C) ventilation heterogeneity. (n = 3-7 animals/genotype, linear regression model, estimated marginal means and 95% CIs, * *p*<0.05, \*\**p*<0.01). Key on graph: Baseline = no PA infection; PA-D07 = 7 days post *P. aeruginosa* bead delivery; PA-D14 = 14 days post *P. aeruginosa* bead delivery; PA-D21 = 21 days post *P. aeruginosa* bead delivery; PA-D63 = 63 days post *P. aeruginosa* bead delivery.

**Figure 8:**
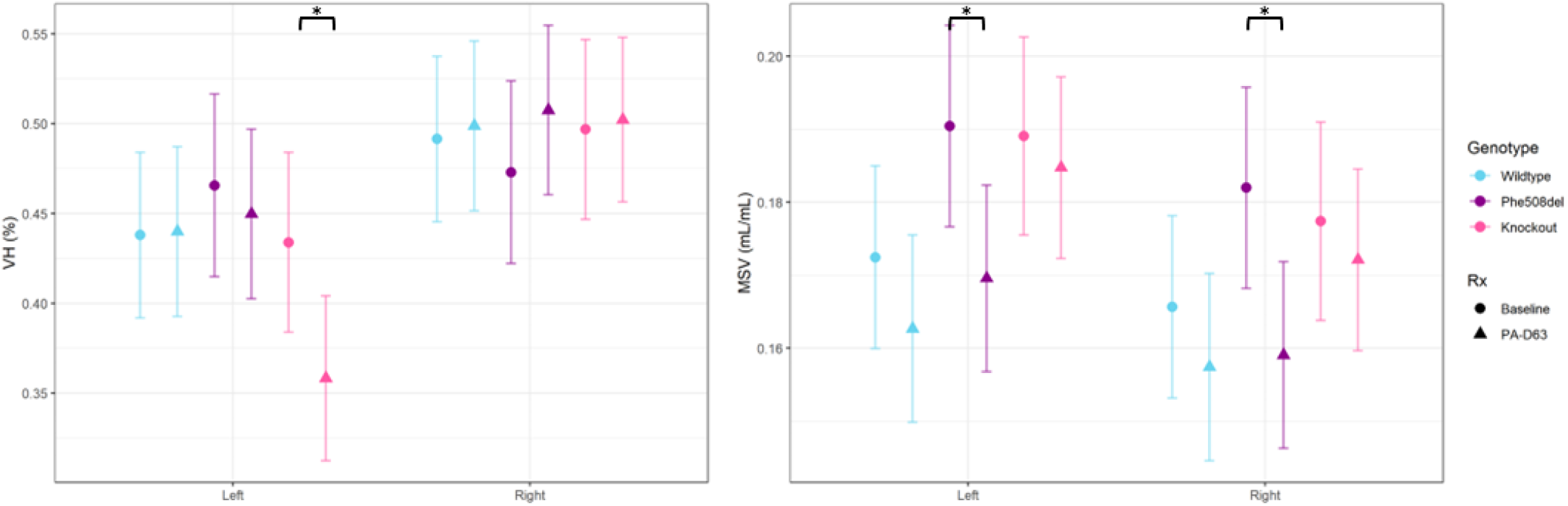
X-ray Velocimetry ventilation of the left and right lungs of wildtype, Phe508del and knockout at baseline and day 63 post *P. aeruginosa* infection. (n = 7-8 animals/genotype, linear regression model, estimated marginal means and 95% CIs, * *p*<0.05). Key on graph: Baseline = no *P. aeruginosa* infection; PA-D63 = 63 days post *P. aeruginosa* bead delivery

To investigate differences in ventilation, the specific ventilation data for each lung was split into left and right components for analysis of the day 63 data. In knockout rats there was a significant decrease in ventilation heterogeneity (VH) of the left lung from baseline to day 63. Significant decrease in mean specific ventilation (MSV) was observed in the left and right lung of the *Phe508del* rats from baseline to day 63 post infection.

## Discussion

Developing a *P. aeruginosa* chronic pulmonary infection model in rodents that accurately mimics what is seen clinically in CF has been a longstanding challenge. Delivery of free *P. aeruginosa* often results in either acute infection leading to sepsis and high mortality, or rapid clearance with no sustained infection. To address this, the technique of embedding bacteria in agar beads was developed, and has since been employed in numerous studies to establish chronic lung infections in rodents [10, 18, 23-29]. The delivery method of bacteria-laden agar beads to the lungs in rodents is often via tracheostomy or orotracheal routes [6]. Both methods result in distribution of *P. aeruginosa* throughout the entire lung.

Few studies have used a CF rodent strain combined with a *P. aeruginosa* isolate embedded in agar to establish a chronic infection [18, 24, 25]. These studies created a whole lung infection, administering higher bacterial loads or volumes, such as 3 × 10^6^ 300 μl in CF rats [18] and 1-2 × 10^6^ in 50 μl in mouse models [24, 25]. These doses are higher than the inoculum used in our study (1.2 × 10^5^ CFU/ 50 μl). A study using CF knockout rats investigated differences between younger and older CF rats, finding that only the older CF knockout rats had a sustained infection up to 28 days, likely due to reduced mucus clearance [18]. The younger CF and wildtype rats were able to clear the infection by day 7. Additionally, a mouse study managed to maintain a stable infection for up to three months using *P. aeruginosa* CF isolate embedded agar beads [24]. These studies also reported morality and significant weight loss associated with their chronic infections, suggesting that the established infection was too severe.

In this study, we aimed to establish a chronic infection localised to the right lung only. Using a miniature bronchoscope, we guided the delivery of the agar beads to the top of the right main bronchus. Due to the upright position of the rat during delivery, the agar beads tended to settle in lower regions of the right lung. At earlier time points (days 7 to 21), histological examination revealed that the upper regions of the right lung and entire left lung remained unaffected. In contrast, the lower right lung exhibited histological changes typically associated with CF lung disease, including acute bronchitis/bronchiolitis and goblet cell hyperplasia. However, by day 63, these changes were observed throughout the right lung, with some effects also appearing in the left side. In people with CF, the upper lung regions, are often the most severely affected, while the rest of the lung may be less involved [30]. This difference is likely due to the upright delivery method used in rats, where the agar beads in a large volume likely settle in the lower lung, which is unlike the humans when bacteria are inhaled and are trapped in the upper airways.

Throughout the study, the overall health of the rats remained stable, evidenced by consistent weight gain, absence of infection-related mortality, and good overall body condition and well-being. The inflammatory response was elevated 7-days post infection, evident by an increased percentage of neutrophils in the BAL fluid from rats of all genotypes. While neutrophil levels in knockout rats declined over the course of the infection, they remained elevated at day 21 when compared to baseline, indicating a sustained inflammatory response. Analysis of immune markers such as TNF-a or IL-1B, could provide further insights to the inflammatory response in this model.

There was no difference in the CFU counts among the genotypes at any time point. However, in wildtype rats, bacterial burden decreased by day 21 to below the initial inoculum and continued to decline up to day 63. In contrast, CF rats (*Phe508del* and knockout) maintained a consistent bacterial load until day 21. By day 63, *Phe508del* rats exhibited a reduction in bacterial burden below the initial inoculum, whereas knockout rats sustained the infection. These findings suggests that CF rats may have impaired ability to clear the infection, consistent with previous studies [18, 31].

Initially, the localised nature of the infection may not have substantially impacted overall (global) lung function, as the lungs have a remarkable ability to compensate; with unaffected areas compensating for the impaired regions [32]. Additionally, histological analysis did not reveal widespread muco-obstruction in the airways, which may explain why typical CF-like changes to lung function were not observed. However, extending the infection to nine weeks appeared to be sufficient to induce tissue changes that significantly affected lung function.

Lung function and mechanics remained largely unchanged during the first 21 days of the infection. By day 63, however, dynamic compliance increased, and elastance decreased across all genotypes, indicating greater lung distensibility. These findings were supported by the pressure-volume loop shifts, suggesting structural lung changes. Histological evidence of emphysema, including alveolar destruction and enlarged airspaces, further aligned with these functional impairments.

Previous studies investigating lung function in a *P. aeruginosa* model or lipopolysaccharide (LPS) derived from *P. aeruginosa* models did not observe significant lung function changes, though these infections were short, lasting less than 7 days [33, 34]. In contrast, a longer study in CF mice using aerosolised LPS administered three times per week for 6 weeks, followed by a 10-week recovery, reported a significant increase in airway resistance [35]. These studies suggest that the short-term infections may not be sufficient to induce the tissue changes needed to affect lung function, whereas longer more sustained infections are more likely to produce detectable functional changes, as seen in this study

XV did not reveal substantial alterations in lung ventilation parameters during the first 21 days. Baseline variability and differing responses to infection between rats may have obscured subtle changes, making it difficult to detect group-level differences. However, at day 63, when XV data were analysed by lung side, changes in MSV were evident in *Phe508del* rats. Knockout rats showed a reduced VH in the left lung, while *Phe508del* rats showed reductions in MSV in both the left and right lungs compared to baseline, indicating a more widespread ventilation impairment by this stage of infection. This decline in MSV along with the histological changes of airway inflammation and structural lung changes, suggests impaired airflow possibly due to airway blockages. A repeated measures study design, where baseline XV measurements are taken and the same animals are tracked over the course of the infection, may help uncover changes. This approach also supports the concept of using XV as a personalised diagnostic tool to monitor lung disease progression in people with CF. However, due to the design of the present study - requiring CFU quantification and histological assessment - we used separate cohorts of rats at each time point and could not longitudinally track individual rats throughout the infection.

A limitation of this study is that we did not include a control group to investigate the impact of sterile bead delivery to the lung. However, previous research has shown that the delivery of sterile agar beads do not cause inflammation like the bacteria-laden beads [18, 27], and we have previously demonstrated the acute effects of sterile agar bead delivery on lung function [14]. Although the infection in this model may be considered relatively mild compared to the other *P. aeruginosa* infection models, it is important to strike a balance between delivering a sufficient bacterial load to the animal to maintain a chronic infection, without giving too much as to cause sepsis and high mortality. Given that the bacterial load and delivery method were well tolerated in this study, future research could explore the use of higher bacterial loads, or repeated bacteria deliveries to increase the severity of the infection. It is also essential to recognize that CF lung disease is progressive and develops over many years with a complex interplay of multiple pathogens. The delivery of *P. aeruginosa* to a relatively sterile lung cannot fully replicate the sustained inflammation, tissue remodelling and tissue damage observed in people with CF. Future models should consider incorporating additional pathogens to better mimic the multi-pathogen nature of CF lung disease.

Another limitation of this study is that the wildtype rats did not fully clear the bacterial infection as expected, instead also developing a chronic disease. This outcome may be due to the model of embedding *P. aeruginosa* into agar beads allowing the bacteria to remain in the lung. As the agar beads break down, there is a sustained and prolonged release of *P. aeruginosa* leading to continuous exposure to the bacteria and the development of chronic disease in the wildtype rats.

This study established a new method for inducing a localised, chronic *P. aeruginosa* infection in CF rats, offering a valuable tool for CF research. The precise delivery of bacterial laden beads using a miniature bronchoscope provides a controlled and reproducible infection that persists for up to nine weeks, with minimal impacts on overall animal health. This infection model could be useful in studies investigating approaches for treating *P. aeruginosa* bacterial infection, particularly targeted therapy due to the localised nature of the infection.

## Acknowledgements

The authors acknowledge the facilities and scientific and technical assistance of the National Imaging Facility, a National Collaborative Research Infrastructure Strategy (NCRIS) capability, at the Large Animal Research and Imaging Facility, South Australian Health and Medical Research Institute. The authors acknowledge Wick Lakshantha and Ryan O’Hare Doig for operating the Permetium small animal scanner. Thanks to Prof Jian Li (Monash University, Australia) for supplying the *P. aeruginosa* strain and A/Prof Susan Birket (University of Alabama at Birmingham, USA) for providing advice related to the *P. aeruginosa* agar bead production method. This project built on experiments M13447, M14029 and M14774 performed on the Imaging and Medical Beamline at the Australian Synchrotron, and the authors thank Kaye Morgan, Marcus Kitchen, and Chantelle Carpentieri for their input.

## Funding Information

Studies were supported by Medical Research Future Fund Grant RFRHPSI000013 and Cystic Fibrosis Foundation Grant DONNEL21GO.

## Authors’ Contributions

NR: Conceptualisation, Methodology, Formal analysis, Data Curation, Investigation, Visualisation, Writing - Original Draft

BB, PC, AM, NRP: Conceptualisation, Methodology, Investigation, Data Curation, Writing – Review & Editing RS: Formal analysis, Software.

NE, KN, JF, ML, JL: Formal analysis, Writing – Review & Editing

DP: Conceptualisation, Resources, Supervision, Writing - Original Draft

MD: Conceptualisation, Data Curation, Formal analysis, Funding acquisition, Investigation, Methodology, Supervision, Visualization, Writing - Original Draft

## Author Disclosure

MD and DP were involved in the research development and validation of the XV technology and have personally purchased shares in 4DMedical.

NE and KN employed by 4DMedical.

## Data Availability

The data generated in this study is available from the authors on reasonable request.

